# Cardiology researchers’ practices and perceived barriers to open science: an international survey

**DOI:** 10.1101/2023.06.29.546350

**Authors:** Kelly D Cobey, Mohsen Alayche, Sara Saba, Nana Yaa Barnes, Sanam Ebrahimzadeh, Emilio Alarcon, Benjamin Hibbert, David Moher

**Affiliations:** University of Ottawa Heart Institute, Ottawa, Canada; School of Epidemiology and Public Health, University of Ottawa, Ottawa, Canada; Department of Medicine, Faculty of Medicine, University of Ottawa; Biopharmaceutical Science, University of Ottawa, Canada; Health Sciences, University of Ottawa, Canada; Centre for Journalology, Ottawa Hospital Research Institute, Ottawa, Canada; Department of Biochemistry, Microbiology and Immunology, Faculty of Medicine, University of Ottawa, Canada; Division of Cardiac Surgery, University of Ottawa Heart Institute, Ottawa, Canada

## Abstract

**Background:** Open science is a movement and set of practices to conduct research more transparently. Implementing open science will significantly improve public access and supports equity. It also has the potential to foster innovation and reduce duplication through data and materials sharing. Here, we survey an international group of researchers publishing in cardiovascular journals regarding their perceptions and practices related to open science.

**Methods:** We identified the top 100 “Cardiology and Cardiovascular Medicine” subject category journals from the SCImago journal ranking platform. This is a publicly available portal that draws from Scopus. We then extracted the corresponding author’s name and e-mail from all articles published in these journals between March 1, 2021 - March 1, 2022. Participants were sent a purpose-built survey about open science. The survey contained primarily multiple choice and scale-based questions for which we report count data and percentages. For the few text-based responses we conducted thematic content analysis.

**Results:** 198 participants responded to our survey. Participants had a mean response of 6.8 (N=197, SD=1.8) on a 9-point scale with endpoints, not at all familiar (1) and extremely familiar (9), when indicating how familiar they were with open science. When asked about where they obtained open science training, most participants indicated this was done on the job self-initiated while conducting research (n=103, 52%), or that they had no formal training with respect to open science (n=72, 36%). More than half of the participants indicated they would benefit from practical support from their institution on how to perform open science practices (N=106, 54%). A diversity of barriers to each of the open science practices presented to participants were acknowledged. Participants indicated that funding was the most essential incentive to adopt open science.

**Discussion:** It is clear that policy alone will not lead to the effective implementation of open science. This survey serves as a baseline for the cardiovascular research community’s open science performance and perception and can be used to inform future interventions and monitoring.

## Introduction

Open science is a movement to make the research lifecycle accessible to all – including practices such as open access publishing, data and code sharing, and open (source) materials sharing. There is growing momentum globally to see open science practices more firmly embedded into the research ecosystem, with several jurisdictions having introduced policies and roadmaps to foster effective implementation^1–5^. Previous research suggests that up to 85% of research conducted is wasted^6^, and that the scientific system is fraught with issues including publication bias, inadequate reporting, and lack of reproducibility^7, 8^. Implementing open science could reduce unnecessary duplication of research, thus saving time and money. Further, open science enhances transparency by making the various components of the research life cycle accessible thereby reducing bias but also driving innovation as others can use and adapt study data and materials. Open science also helps to support equity by reducing barriers in access to information. In medicine this means that researchers and the public alike do not face barriers in accessing health information. Despite the growing impetus to implement open science globally, no country or discipline has achieved widespread adoption. There are several real and perceived challenges transitioning from our current norm of ‘closed research’. Issues including how to effectively create behavior change to promote open science activities, how to train researchers on the formal practices involved with open science, and how to reconcile openness with intellectual property, have all been raised as potential challenges.

The current study investigates cardiology researcher’s perceptions and practices related to open science. A recent cross-sectional study examining 232 publications in cardiology journals found that 96.6% (N= 224/232) of papers audited did not have publicly available data, while 229/232 (98.7%) did not provide their analysis scripts, and 98.3% (228/232) did not refer to an accessible study protocol^9^. Other related research has shown that study design and reporting elements to reduce bias in preclinical cardiology studies (e.g., blinding, randomization) are not common within the field and that this issue has persisted without much improvement year on year^10^. Collectively, this work suggests that open science practices and related reporting and design best practices are not normative within cardiology. Given this reality, it is little wonder why concerns about reproducibility in the field persist^11–13^. We know of no study to date that has surveyed cardiology researchers’ perceptions of open science. This is regrettable as knowledge of researchers’ perceptions of open science, and of barriers and facilitators to achieving openness, is essential to understand how to support the community to implement open science more fully. Other disciplines including social science^14^, economics^15^, and psychology^16–18^ have conducted large-scale surveys of their researchers to determine the state of open science in their community. Such surveys can be used as a starting point to develop interventions to implement open science more effectively, but they can also serve to monitor open science over time with additional surveys that can be compared longitudinally. We used a cross-sectional online survey, sent to a randomly selected sample of corresponding authors of recent publications in well-known cardiology journals, to measure perceptions and practices related to open science. The study is descriptive, and we have no hypotheses. The survey is the first in a program of research we are leading to target implementation of open science in cardiology^19^.

## Methods

### Transparency statement

This study received ethical approval from Ottawa Health Sciences Research Ethics Board. All study materials and data are available on the Open Science Framework^20^ along with the study registration: https://osf.io/v42u8/ ^21^.

### Study Design

We conducted a cross-sectional online survey sent to a randomly selected sample of corresponding authors of publications in cardiology journals.

### Sampling framework

We identified the top 100 “Cardiology and Cardiovascular Medicine” subject category journals from the SCImago journal ranking platform. This is a publicly available portal that draws from Scopus. We then extracted the corresponding author’s name and e-mail from the all articles published in these journals between March 1, 2021 - March 1, 2022. We included authors of all article types. For full details on our approach to extracting author emails please see Appendix 1. This is a convenience sample, because the work is descriptive and we are not conducting any inferential tests, we did not conduct a power analysis.

### Participant Recruitment

This closed survey was sent to researchers who we identified through our sampling framework. Potential participants received an email including an approved recruitment script that explained the study’s aim and invited them to complete our anonymous online survey. Involvement in the survey served as implied consent. There was no incentive to take part in the survey.

We used Mail Merge software to send emails to the authors in our sample. We sent three reminder emails to participants at weekly intervals from the original invitation to encourage responses and closed the survey four weeks after the initial invitation was received. After de-duplication of repeated emails, we sent our recruitment script to a total of 9594 researchers. We received 844 bounce backs, meaning a total our sample was 8,750 researchers.

### Survey

The full survey is available in Appendix 2. Participants were asked six demographic questions (e.g., gender, age). Following this, they responded to 3 questions about their research expertise and role. Then, participants were asked to indicate their familiarity with open science.

Subsequent questions asked about participants’ training related to open science. Participants were presented with definitions of open access publishing, preprints, data sharing, materials sharing, protocol registration, reporting guidelines, and patient engagement, and asked whether they had experience performing the practice and what barriers they face to so. Most of the questions were multiple choice and participants could navigate through a back button. Prior to completion, the survey was pilot tested by two cardiology researchers for clarity and format, with their feedback integrated into the design. We estimate that completing the survey took 10 minutes. Participants had the option of skipping any questions that they do not wish to answer.

### Data Analysis

Data analysis was conducted using Excel. We report basic descriptive statistics (e.g., counts, percentages). Rather than conducting Chi Square Crosstabs tests to test for group differences in responses (e.g., considering gender, career stage, cardiology sub-discipline) as per our protocol, we have provided descriptive tables of these group differences given modest group sizes. For text-based responses, two members of the research team conducted a thematic content analysis. To do so, each researcher coded responses separately. Following a discussion and iterative updates to obtain a consensus on the codes, they were conceptually organized into topic areas and defined and explained in tables for reporting.

## Results

### Demographics

A total of 198 individuals completed the survey (response rate 2.3%). Participants tended to be male (N=153, 77%) and based in North America (N=97, 49%). Most participants reported to be faculty members/primary investigators (N=152, 77%), primarily working in clinical research (N=127, 64%), and that cardiovascular research was their main research area (N=178, 90%). For complete demographics, please see Table 1.

**Table 1.** Participant demographics

| Item | Response options | N | % |
| --- | --- | --- | --- |
| Which describes you best? | Male | 153 | 77% |
|  | Female | 43 | 22% |
|  | Non-binary | 1 | 1% |
|  | Blank | 1 | 1% |
| What is your age group? | 18-24 | 1 | 1% |
|  | 25-34 | 24 | 12% |
|  | 35-44 | 65 | 33% |
|  | 45-54 | 50 | 25% |
|  | 55-64 | 33 | 17% |
|  | 65 or older | 24 | 12% |
|  | Blank | 1 | 1% |
| Do you self-identify as having a disability? | Yes | 3 | 2% |
|  | No | 192 | 97% |
|  | Prefer not to say | 2 | 1% |
|  | Blank | 1 | 1% |
| Do you identify as being part of a visible minority group? | Yes | 25 | 13% |
|  | No | 165 | 83% |
|  | Prefer not to say | 4 | 2% |
|  | Blank | 4 | 2% |
| Do you currently identify as a caregiver (i.e., parenting kids under 18, caring for elderly relatives)? | Yes | 95 | 48% |
|  | No | 98 | 49% |
|  | Prefer not to say | 1 | 1% |
|  | Blank | 4 | 2% |
| Where are you located? | North America | 97 | 49% |
|  | Asia | 11 | 6% |
|  | South America | 7 | 4% |
|  | Europe | 72 | 36% |
|  | Australasia | 9 | 5% |
|  | Africa | 1 | 1% |
|  | Blank | 1 | 1% |
| Which of the following describes you best? | Faculty member/PI | 152 | 77% |
|  | Postdoctoral fellow | 13 | 7% |
|  | Graduate student | 9 | 5% |
|  | Scientist in third sector (E.g., NGO, non-profit) | 2 | 1% |
|  | Research support staff (E.g., research manager, research associate, technician) | 6 | 3% |
|  | Scientist in industry | 3 | 2% |
|  | Government scientist | 6 | 3% |
|  | Other | 6 | 3% |
|  | Blank | 1 | 1% |
| Which of the following best describes your primary research area? | Clinical research | 127 | 64% |
|  | Health systems research | 13 | 7% |
|  | Epidemiological research | 15 | 8% |
|  | Preclinical research – in vivo | 23 | 12% |
|  | Methods research | 6 | 3% |
|  | Preclinical research – in vitro | 6 | 3% |
|  | Other | 7 | 4% |
|  | Blank | 1 | 1% |
| Is cardiovascular research your main area of research? | Yes | 178 | 90% |
|  | No | 20 | 10% |

### Open science familiarity, training, and incentivization

Participants had a mean response of 6.8 (N=197, SD=1.8) on a 9-point scale with endpoints, not at all familiar (1) and extremely familiar(9), when indicating how familiar they were with open science. When asked about where they obtained open science training most participants indicated this was done on the job self-initiated while conducting research (n=103, 52%), or that they had no formal training with respect to open science (n=72, 36%). Participants indicated their top format preference for training related to open science would be a website of resources. Additional funding to perform open science practices was the top incentive listed by participants to encourage them to apply more open science practices (N=154, 78%). More than half of participants indicated they would benefit from practical support from their institution on how perform open science practices (N=106, 54%). Funders and research institutions were the top indicated stakeholders in terms of which has the most ability to create policies that result in successful uptake of open science. For complete results please see Table 2.

**Table 2.** Open science education and motivation

| Item | Responses | Yes (N) | % |
| --- | --- | --- | --- |
| Most of my training with respect to open science has been learned: | On the job self-initiated while conducting research | 103 | 52% |
|  | I have no formal training with respect to open science | 72 | 36% |
|  | Through mentorship directly from my supervisor and/or peers | 11 | 6% |
|  | Via formal coursework/workshops instructing about open science | 8 | 4% |
|  | Other | 3 | 2% |
|  | Blank | 1 | 1% |
| Item | Rank | Responses |  |
| If you were to engage in training related to open science, which format of training would be your preference? (Top 3) | 1 | A website of resources |  |
|  | 2 | An online webinar/recording |  |
|  | 3 | A short online course of six sessions (asynchronous) |  |
| Item | Responses | Yes (N) | % |
| Which of the following incentives would result in you applying more open science practices? | Clearer communication about why open science is valuable for research | 77 | 39% |
|  | Practical support from my institution to conduct open science | 106 | 54% |
|  | Additional funding to perform open science practices | 154 | 78% |
|  | Additional training on how to perform open science practices | 61 | 31% |
|  | Having a staff trained on open science practices | 53 | 27% |
|  | A way to get recognized for my performance of open science practices when I am being hired/promoted/tenured] | 58 | 29% |
|  | Other | 9 | 5% |
| Item | Rank | Responses |  |
| Rank order the stakeholders below in terms of which you feel has the most significant impact on creating policies that result in successful uptake of open science. | 1 | Funders |  |
|  | 2 | Research institutions |  |
|  | 3 | Scholarly journals |  |

Free text responses to the item asking about the best ways to promote open science were coded into 25 unique codes. These codes were then thematically grouped which resulted in 7 categories: 1) Finances, 2) Incentives, 3) Policy and guidance, 4) Support, 5) Culture change, 6) Perceived concerns, and 7) Other. Illustrative examples of each theme are provided in Table 3.

**Table 3.** Thematic analysis of best ways to promote open science

| Theme | Code | N (137) | % | Example |
| --- | --- | --- | --- | --- |
| Finances | Provide funding | 30 | 21.9 | “Provide funding at the institutional level to support it” |
|  | Make it affordable | 11 | 8.0 | “make it much less expensive” |
|  | Don't charge fees to researchers | 7 | 5.1 | “Don't charge scientists to do it” |
| Incentives and rewards | Incentivize open science practices | 4 | 2.9 | “The incentive has to be correct, but one can not expect to give data away (as some initiatives almost look like).” |
|  | Use novel metrics | 4 | 2.9 | “Use novel metrics (Altmetrics) instead of impact factors.” |
|  | Support broader types of publication models | 3 | 2.2 | “Creation of open repository for scientific publications, and promotion of the repository through media.” |
|  | Show value to researchers | 6 | 4.4 | “Have research funders and universities promote it more, explain what it entails, and the benefits to the researcher and science...” |
|  | Provide recognition for open science activities | 3 | 2.2 | “Ensure that those who have spent years dedicated to generating the data continue to receive primary academic/ institutional credit for the work.” |
| Policy and process changes | Create mandates | 11 | 8.0 | “Embed it as a requirement in nationally funded research” |
|  | Improve peer review | 2 | 1.5 | “Make it peer-reviewed.” |
|  | Disseminate guidelines | 2 | 1.5 | “Emphasize the importance of open science in society guidelines..” |
| Support | Facilitate the process | 7 | 5.1 | “Institutional arrangements with publishers to ensure free Open Access publication in hybrid form” |
|  | Provide training | 12 | 8.8 | “undergraduate training” |
|  | Knowledge sharing of practices | 2 | 1.5 | “give example on how this will give access to data and protocols develop elsewhere..” |
| Culture change | Change journal culture | 8 | 5.8 | “Change the journal culture.” |
|  | Consider geographic realities | 2 | 1.5 | “...researchers from LMICS would simply publish under a subscription model..” |
|  | Research culture change | 4 | 2.9 | “Making data available would be the single step. However, the converse is that data is a scientists 'currency' for advancing their own research and exposes one to having ideas stolen..” |
| Perceived concerns | Address low quality journals | 4 | 2.9 | “Open science is hampered by all the really bad journals promoting open-access paid articles.” |
|  | Alleviate concern of misinformation | 3 | 2.2 | “Alleviate concerns that data will not be misinterpreted, and reports will not have incorrect conclusions” |
|  | Ensure open science is rigorous | 3 | 2.2 | “I am not an unconditional fan of opens science as it is being promoted presently and therefore do not think it is appropriate to promote it without further consideration as it is currently done” |
| Other | Other | 9 | 6.6 | “The biggest barrier, by far, is the risk associated with open data from institutional review boards and the privacy act.” |

### Open science performance

Most participants reported having experience publishing an article open access (N=168, 85%) and using a reporting guideline (N=123, 62%). Roughly half of researchers reported that they had experience registering a study protocol (N=106, 54%) or engaging patients or members of the public in research (N=96, 48%). Fewer researchers reported experience sharing study materials (n=54, 27%), making a preprint (N=49, 25%), or sharing study data (N=48, 24%). Please see Table 4.

**Table 4.** Open science performance

| Item | Responses | N | % |
| --- | --- | --- | --- |
| **In that past 12 months have you published an article ‘open access’?** | Yes | 168 | 85% |
|  | No | 28 | 14% |
|  | I don't know | 1 | 1% |
|  | I have not published a research paper in the past 12 months | 1 | 1% |
| **In the past 12 months have you made a preprint prior to publishing an article?** | Yes | 49 | 25% |
|  | No | 139 | 70% |
|  | I don't know | 7 | 4% |
|  | Blank | 3 | 2% |
| **In that past 12 months have you shared the raw data (all data necessary for reproducing the research) underpinning a study at the time of publication?** | Yes | 48 | 24% |
|  | No | 147 | 74% |
|  | I don't know | 1 | 1% |
|  | Blank | 2 | 1% |
| **In the past 12 months have you shared the study materials underpinning a study at the time of publication?** | Yes | 54 | 27% |
|  | No | 131 | 66% |
|  | I don't know | 9 | 5% |
|  | Blank | 4 | 2% |
| **In that past 12 months have you registered a study protocol for any research project you are working on?** | Yes | 106 | 54% |
|  | No | 82 | 41% |
|  | I don't know | 1 | 1% |
|  | I have not initiated a research study in the past 12 months | 8 | 4% |
|  | Blank | 1 | 1% |
| **In that past 12 months have you explicitly used and referenced a reporting guideline checklist in any research report you have published?** | Yes | 123 | 62% |
|  | No | 66 | 33% |
|  | I don't know | 6 | 3% |
|  | I have not published a research paper in the past 12 months | 1 | 1% |
|  | Blank | 2 | 1% |
| **In that past 12 months have you engaged patients or members of the public in any research you have conducted?** | Yes | 96 | 48% |
|  | No | 94 | 47% |
|  | I don't know | 3 | 2% |
|  | I have not conducted a research project in the past 12 months | 3 | 2% |
|  | Blank | 2 | 1% |

### Barriers to open science

A diversity of barriers to each of the open science practices presented to participants were acknowledged. For the complete results, please see Table 5. When we asked participants about barriers to publishing their work open access, the top barrier identified was funding to support open access article processing charges (N=149, 75%). A fifth of participants also indicated that they did not perceive their institution valued open access publishing (N=40, 20%). When asked about the barriers to creating a preprint almost half of participants indicated that they felt there were potential harms associated with work that has not been peer reviewed (N=91, 46%). Other key barriers included that participants worried that making a preprint would reduce their chances of the work being accepted at a peer reviewed journal (N=73, 37%), and that they did not see the benefit of making a preprint (N=71, 36%).

**Table 5.** Barriers to open science

| Barrier | N | % |
| --- | --- | --- |
| Open access publishing |  |  |
| The journals in my area don't use an open access publishing model | 15 | 8% |
| I don't know how to self-archive a paper to make it open access | 24 | 12% |
| I don't see the benefit of making an article open access | 15 | 8% |
| I don't think my institution values me doing this | 40 | 20% |
| I don't have funding to support the article processing charges that are common at open access journals | 149 | 75% |
| I do not perceive any of the above as issues to publish open access | 20 | 10% |
| Other | 21 | 11% |
| **Preprints** |  |  |
| I don't really know how to make a preprint | 52 | 26% |
| I don't have time to make preprints | 35 | 18% |
| I worry making a preprint will reduce my chances of the work being accepted at a peer reviewed journal | 73 | 37% |
| I don't see the benefit in making a preprint | 71 | 36% |
| I don't think my institution values me making a preprint | 55 | 28% |
| I think there are potential harms associated with sharing work that has not been peer reviewed | 91 | 46% |
| My institution has an internal process for posting preprints that makes the process very time consuming | 7 | 4% |
| Other | 13 | 7% |
| **Data sharing** |  |  |
| I don't know how to prepare my data appropriately for sharing | 45 | 23% |
| I don't have time to prepare my data for sharing | 59 | 30% |
| I don't know where to share my data | 44 | 22% |
| I don't feel I will get recognition for sharing my data | 65 | 33% |
| My institutional ethics board will not allow me to share my data | 58 | 29% |
| My research consent form specifies I will not share the data | 48 | 24% |
| I am concerned about patient privacy if I share my data | 72 | 36% |
| Concerns about intellectual property control | 94 | 47% |
| Concerns about being scooped | 56 | 28% |
| Concerns about unintended use of secondary data | 88 | 44% |
| Concerns about misinterpretation of the data | 75 | 38% |
| Concerns others may discover errors in the data | 15 | 8% |
| Other | 23 | 12% |
| **Materials sharing** |  |  |
| I don't know how to prepare my study materials for sharing | 38 | 19% |
| I don't have time to prepare my study materials for sharing | 43 | 22% |
| I don't know where to share my study materials | 46 | 23% |
| I don't think there is value for others in me sharing my study materials | 21 | 11% |
| There is no appropriate infrastructure available for me to share my study materials | 49 | 25% |
| The costs to share my study materials are a barrier | 41 | 21% |
| I don't feel I will get recognition for sharing my study materials | 51 | 26% |
| My institutional ethics board will not allow me to share my study materials | 26 | 13% |
| I am concerned about patient privacy if I share my study materials | 35 | 18% |
| Concerns about intellectual property control | 63 | 32% |
| Concerns about being scooped | 35 | 18% |
| Concerns about unintended use of materials | 54 | 27% |
| Concerns about misinterpretation of the materials | 40 | 20% |
| There aren't any trained staff to guide me about it | 21 | 11% |
| Other | 20 | 10% |
| **Protocol registration** |  |  |
| I don't know how to create a study registration | 13 | 7% |
| I don't know what platform to use to register my study | 24 | 12% |
| I don't have time to register my studies | 33 | 17% |
| I don't feel I will get recognition for taking the time to register my studies | 24 | 12% |
| I don't think that my institution prioritizes study registration | 24 | 12% |
| I worry that I will be scooped if I share my study plan before publishing results | 22 | 11% |
| I don't think there is value for others in me registering my studies | 18 | 9% |
| I don't think my research area lends itself well to registering protocols | 20 | 10% |
| Other | 23 | 12% |
| **Reporting guidelines** |  |  |
| I don't know where to find the relevant reporting guideline | 23 | 12% |
| I don't know how to use reporting guidelines | 17 | 9% |
| I don't have time to use reporting guidelines | 23 | 12% |
| I don't see the value in using reporting guidelines | 21 | 11% |
| I don't feel I will get recognition for taking the time to user reporting guidelines | 27 | 14% |
| I don't think my institution prioritizes the use of reporting guidelines | 17 | 9% |
| Other | 35 | 18% |
| **Patient and public involvement** |  |  |
| I don't know how to identify patients/public members to contribute | 33 | 17% |
| I don't know how to incorporate patients/public members in my research | 48 | 24% |
| I don't have time to incorporate patients/public members in my research | 30 | 15% |
| I don't see the value in incorporating patients/public members in my research | 20 | 10% |
| I don't feel I will get recognition for taking the time to incorporate patients/public members in my research | 28 | 14% |
| I don't think my institution prioritizes incorporating patients/public members in my research | 23 | 12% |
| Other | 35 | 18% |

When asked about barriers to sharing study data openly almost half of participants indicated they had concerns about intellectual property control (N=94, 47%). Other key barriers were participants’ concern about unintended use of secondary data (N=88, 44%) and concerns about misinterpretation of the data (N=75, 38%). Participants also raised concerns about intellectual property (N=63, 32%) and concerns about unintended use (N=54, 27%) as key barriers to materials sharing.

When asked about barriers to protocol registration, use of reporting guidelines, and patient and public involvement, no overwhelming majority emerged for a particular item. The top barriers noted were that participants don’t have time to register studies (N=33, 17%), participants do not feel they get recognition for taking time to use reporting guidelines (N=27, 14%), and that participants don’t know how to incorporate patients/public members in their research (N=48, 24%).

## Discussion

We report the results of a survey of the cardiovascular community’s perceptions and experiences with open science. Our results compliment previous discipline specific efforts in social science^14^, economics^15^, and psychology^16–18^ which have begun to provide data about the unique challenges of implementing and fostering open science in particular disciplines. Given the prevalence of cardiovascular diseases globally efforts to embed open science within this discipline have great potential to increase the useability and integrity of research in this area and ultimately to have a positive downstream impact on patient treatment and prevention of cardiovascular disease. Our findings provide an important baseline that can be used to track progress in open science implementation over time.

We found that most participants had either no formal open science training or had obtained training on the job on their own. This suggests that most researchers in the cardiovascular research community are figuring out open science as they encounter it, rather than any sort of cohesive community approach to implementation. This is especially concerning since participants also indicated that clearer communication about why open science is valuable for research would incentivize them to implement open science. Together, it suggests that participants need to be better and more systematically supported.

Participants indicated funding was the most essential incentive to adopt open science. The need for funding was also reflected in the thematic analysis of what is needed to best promote open science, where three codes related to financing open science. While much of this discussion focused on for practicing open access publishing, which typically is associated with an article processing charge, the call for funding to hire personnel to carry out open science activities (e.g., data management) was also made. The second most important incentive among participants was support from their institutions to conduct open science. Such support may take the form of toolkits or training, but support in the form of personal was again noted as valuable.

Participants indicated that they perceived funders had the most significant impact on creating policies that result in successful uptake of open science. This suggests that the community feels the need to respond to funder policies; however, when we examine the rates of self-reported performance of open science practices that are commonly mandated, we see a gap in performance. Overall, rates of self-reported performance of open science practices were limited. Eighty five percent of respondents indicated they had published an open access article in the past year, while 62% indicated they had used a reporting guideline checklist. Mandates for both of these practices at the funder and journal level, respectively, are the norm. e.g.^22–24^. Funders implementing audit of the open science practices they mandate may help to ensure these and other practices are being implemented optimally.

About half of participants additionally indicated that they had registered a study protocol. This is interesting, given that mandates only exist for clinical trials^25, 26^, suggesting that the broader recognition of publication bias and selective outcome reporting^27, 28^ may be leading to more general study registration albeit still at sub-optimal levels. Almost half of participants indicated that they had engaged patients in research, yet 24% said they did not know how to incorporate patients/public members suggesting that there remains a need for awareness raising. Rates of the remaining open science practices were comparatively low, suggesting that they are even less embedded into the ethos of the average cardiovascular researcher.

When considering barriers to implementing each of the various open science practices, participants noted concerns that represent a lack of understanding or expertise on open science topics. For example, 37% worried that making a preprint would harm there chances of later publishing, despite the fact that preprints are nearly uniformly accepted at biomedical journals and that there are tools to check this^29^. Another example is the concern about intellectual property control when sharing data and materials, which may suggest a need to outreach on how open science is compatible with a pathway for collaborative R&D^30^.

As a next step the barriers common to participants for each of the open science practices examined can be used to develop interventions to improve performance on the practice. Our survey will serve as a baseline for the community to tracks its progress on implementing open science but also on tracking what barriers persist and change over time to be responsive to the needs of the community.

While our study benefited from a broad sampling strategy and diverse participation, it has several limitations. Given the increasing mandates for open science practices it is possible that some participants did not feel comfortable providing answers that presented themselves in an unfavorable way. Survey question answer responses, particularly those presented when asking about barriers to the various open science practices, may not have fully represented participants views and may potentially have been interpreted differently by different participants. Finally, while the random sampling strategy we undertook allowed us to sample a diverse range of cardiovascular researchers, it is possible that those that responded to our survey, which was not incentivized in anyway, may differ in some way from those who opted not to respond. This selection bias limitation is a common weakness of survey designs, and we have no way of knowing if our sample matched the population of potential participants, we invited to complete the survey.

We hope these findings will provide valuable data to discuss as a cardiovascular research community and as we endeavor to bring the community together to contribute to a roadmap to implanting open science^19^.

## Acknowledgements

We are grateful to Phoebe Nguyen for her helpful discussion regarding our sampling approach.

## Author contributions

Conceptualization: KDC; Methodology: All authors; Writing-first draft: KDC, SS; Writing-revising and editing: All authors; Supervision: KDC.

## Appendix 1

### Journal Search

We will obtain a list the top 100 ranked “Cardiology and Cardiovascular Medicine” subject category journals from the SCImago journal ranking platform.

### Article Retrieval

We will search for all articles published in each journal using the search strategy *1234567.jc.* where “1234567” is the NLM ID of the journal. Where NLM ID is not available, we will use the syntax “Name of journal”.nj.

We will run search for each journal separately. After each search, we will sort the results by Entry date (descending) and export all publications between March 2021-March 2022.

### Email Retrieval

All retrieved articles will be re-imported to Zotero to retrieve PMID numbers. The list of PMID numbers will be exported as an .csv file and input into an R script (built based on the easyPubMed package) to retrieve the authors’ name, affiliation institutions and email addresses. We will then de-duplicate emails.

## Appendix 2: Study survey

### Demographics

1. Which describes you best?
  - Female
  - Male
  - Non-binary
  - Prefer to self-describe ______
  - Prefer not to say
2. What is your age group?
  - 18-24
  - 25-34
  - 35-44
  - 45-54
  - 55-64
  - 65 or older
  - Prefer not to say
3. Do you identify as being part of a visible minority group?
  - Yes
  - No
  - Prefer not to say
4. Do you self-identify as having a disability?
  - Yes
  - No
  - Prefer not to say
5. Do you currently identify as a caregiver (i.e., parenting kids under 18, caring for elderly relatives)?
  - Yes
  - No
  - Prefer not to say
6. Where are you located?
  - Africa
  - Asia
  - Australasia
  - Europe
  - North America
  - South America
  **Research demographics**
7. Which of the following describes you best?
  - Graduate student
  - Postdoctoral fellow
  - Faculty member/PI
  - Research support staff (E.g., research manager, research associate, technician)
  - Scientist in industry
  - Scientist in third sector (E.g., NGO, non-profit)
  - Government scientist
  - Other, please specify
8. Which of the following best describes your primary research area?
  - Clinical research
  - Preclinical research – in vivo
  - Preclinical research – in vitro
  - Health systems research
  - Methods research
  - Epidemiological research
  - Other, please specify
9. Is cardiovascular research your main area of research? Yes No

### Open Science practice and experience

“Open science is an umbrella term that reflects the idea that scientific knowledge of all kinds, where appropriate, should be openly accessible, transparent, rigorous, reproducible, replicable, accumulative and inclusive, all which are considered fundamental features of the scientific endeavour. Open science consists of principles and behaviours that promote transparent, credible, reproducible and accessible science. Open science has six major aspects: open data, open methodology, open source, open access, open peer review and open educational resources.” (<u>Source</u>: FORRT Glossary)

1. How familiar are you with the concept of open science overall? (1-9-point scale; Not at all familiar – Moderately familiar – Extremely familiar)
2. *Note: There was an item in the protocol additional to the general item above asking participants to respond to how familiar they are with each of the below open science practices, but due to a technical glitch in survey development, it was not presented to all participants*

3. Most of my training with respect to open science has been learned:

| Item | Definition |
| --- | --- |
| Open access | “Free availability of scholarship on the public internet, permitting any users to read, download, copy, distribute, print, search, or link to the full texts of these research articles, crawl them for indexing, pass them as data to software, or use them for any other lawful purpose, without financial, legal, or technical barriers other than those inseparable from gaining access to the internet itself” (Boai, 2002). |
| Preprints | A publicly available version of any type of scientific manuscript/research output preceding formal publication, considered a form of Green Open Access. Preprints are usually hosted on a repository (e.g. arXiv) that facilitates dissemination by sharing research results more quickly than through traditional publication. Preprint repositories typically provide persistent identifiers (e.g. DOIs) to preprints. Preprints can be published at any point during the research cycle, but are most commonly published upon submission (i.e., before peer-review). Accepted and peer-reviewed versions of articles are also often uploaded to preprint servers, and are called postprints. |
| Open data | Open data refers to data that is freely available and readily accessible for use by others without restriction, “Open data and content can be freely used, modified, and shared by anyone for any purpose” (https://opendefinition.org/). Open data are subject to the requirement to attribute and share alike, thus it is important to consider appropriate Open Licenses. Sensitive or time-sensitive datasets can be embargoed or shared with more selective access options to ensure data integrity is upheld. |
| Open materials | Author’s public sharing of materials that were used in a study, “such as survey items, stimulus materials, and experiment programs” (Kidwell et al., 2016, p. 3). Digitally-shareable materials are posted on open access repositories, which makes them publicly available and accessible. Depending on licensing, the material can be reused by other authors for their own studies. Components that are not digitally-shareable (e.g. biological materials, equipment) must be described in sufficient detail to allow reproducibility. |
| Registration | The practice of publishing the plan for a study, including research questions/hypotheses, research design, data analysis before the data has been collected or examined. It is also possible to preregister secondary data analyses (Merten & Krypotos, 2019). A preregistration document is time-stamped and typically registered with an independent party (e.g., a repository) so that it can be publicly shared with others (possibly after an embargo period). Preregistration provides a transparent documentation of what was planned at a certain time point, and allows third parties to assess what changes may have occurred afterwards. The more detailed a preregistration is, the better third parties can assess these changes and |
|  | with that the validity of the performed analyses. Preregistration aims to clearly distinguish confirmatory from exploratory research. |
| Reporting guidelines | A reporting guideline is a “checklist, flow diagram, or structured text to guide authors in reporting a specific type of research, developed using explicit methodology.” (EQUATOR Network, n.d.). Reporting guidelines provide the minimum guidance required to ensure that research findings can be appropriately interpreted, appraised, synthesized and replicated. Their use often differs per scientific journal or publisher. |
| Patient and public involvement | Active research collaboration with the population of interest, as opposed to conducting research “about” them. Researchers can incorporate the lived experience and expertise of patients and the public at all stages of the research process. For example, patients can help to develop a set of research questions, review the suitability of a study design, approve plain English summaries for grant/ethics applications and dissemination, collect and analyse data, and assist with writing up a project for publication. This is becoming highly recommended and even required by funders (Boivin et al., 2018). |

a. On the job self-initiated while conducting research
b. Via formal coursework/workshops instructing about open science
c. Through mentorship directly from my supervisor and/or peers
d. Other, please specify:
e. I have no formal training with respect to open science
4. If you were to engage in training related to open science, which format of training would be your preference? (Rank order)

a. A website of resources
b. A handbook of resources
c. A short online course of six sessions (asynchronous)
d. A short online course of six sessions (live)
e. An online webinar/recording
f. An in-person lecture at my institution
g. An in-person workshop at my institution
5. Which of the following incentives would result in you applying more open science practices? (Check all that apply)

a. Clearer communication about why open science is valuable for research
b. Practical support from my institution to conduct open science (E.g., a person I could turn to ask questions about the practicalities of performing open science)
c. Additional funding to perform open science practices (E.g., funding for open access charges, funding for staff to help prepare data for open sharing)
d. Additional training on how to perform open science practices
e. Having a staff trained on open science practices
f. A way to get recognized for my performance of open science practices when I am being hired/promoted/tenured
g. Other, please specify
6. Rank order the stakeholders below in terms of which you feel has the most significant impact on creating policies that result in successful uptake of open science.
  a. Research institutions
  b. Funders
  c. Scholarly journals
  d. Scholarly societies
7. What do you think is the best way to promote open science? (Free text) **Open access**
8. In that past 12 months have you published an article ‘open access’?
  a. Yes
  b. No
  c. I have not published a research paper in the past 12 months
  d. I don’t know
9. Which of the following are barriers for you with respect to publishing open access? (Check all that apply)

a. The journals in my area don’t use an open access publishing model
b. I don’t know how to self-archive a paper to make it open access
c. I don’t see the benefit of making an article open access
d. I don’t think my institution values me doing this
e. I don’t have funding to support the article processing charges that are common at open access journals
f. I do not perceive any of the above as issues to publish open access
g. Other, please specify **Preprints**
10. In the past 12 months have you made a preprint prior to publishing an article?
  a. Yes
  b. No
  c. I have not published a research paper in the past 12 months
  d. I don’t know
11. Which of the following are barriers for you with respect to creating preprints? (Check all that apply)

a. I don’t really know how to make a preprint
b. I don’t have time to make preprints
c. I worry making a preprint will reduce my chances of the work being accepted at a peer reviewed journal
d. I don’t see the benefit in making a preprint
e. I don’t think my institution values me making a preprint
f. I think there are potential harms associated with sharing work that has not been peer reviewed
g. My institution has an internal process for posting preprints that makes the process very time consuming
h. Other, please specify **Open Data**
12. In that past 12 months have you shared the raw data (all data necessary for reproducing the research) underpinning a study at the time of publication?
  a. Yes
  b. No
  c. I have not published a research paper in the past 12 months
  d. I don’t know
13. Which of the following are barriers for you with respect to you sharing the raw data from your research when publishing? (Check all that apply)
  a. I don’t know how to prepare my data appropriately for sharing
  b. I don’t have time to prepare my data for sharing
  c. I don’t know where to share my data
  d. I don’t feel I will get recognition for sharing my data
  e. My institutional ethics board will not allow me to share my data
  f. My research consent form specifies I will not share the data
  g. I am concerned about patient privacy if I share my data
  h. Concerns about intellectual property control
  i. Concerns about being scooped
  j. Concerns about unintended use of secondary data
  k. Concerns about misinterpretation of the data
  l. Concerns others may discover errors in the data
  m. Other, please specify
  **Open Materials**
14. In that past 12 months have you shared the study materials underpinning a study at the time of publication?
  a. Yes
  b. No
  c. I have not published a research paper in the past 12 months
  d. I don’t know
15. Which of the following are barriers for you with respect to you sharing study materials from your research when publishing? (Check all that apply)
  a. I don’t know how to prepare my study materials for sharing
  b. I don’t have time to prepare my study materials for sharing
  c. I don’t know where to share my study materials
  d. I don’t think there is value for others in me sharing my study materials
  e. There is no appropriate infrastructure available for me to share my study materials
  f. The costs to share my study materials are a barrier
  g. I don’t feel I will get recognition for sharing my study materials
  h. My institutional ethics board will not allow me to share my study materials
  i. I am concerned about patient privacy if I share my study materials
  j. Concerns about intellectual property control
  k. Concerns about being scooped
  l. Concerns about unintended use of materials
  m. Concerns about misinterpretation of the materials
  n. There aren’t any trained staff to guide me about it
  o. Other, please specify
  **Registration**
16. In that past 12 months have you registered a study protocol for any research project you are working on? Yes No I have not initiated a research study in the past 12 months I don’t know
17. Which of the following are barriers for you with respect to registering your study protocol prior to starting the research project? (Check all that apply)
  a. I don’t know how to create a study registration
  b. I don’t know what platform to use to register my study
  c. I don’t have time to register my studies
  d. I don’t feel I will get recognition for taking the time to register my studies
  e. I don’t think that my institution prioritizes study registration
  f. I worry that I will be scooped if I share my study plan before publishing results
  g. I don’t think there is value for others in me registering my studies
  h. I don’t think my research area lends itself well to registering protocols
  i. Other, please specify
  **Reporting guidelines**
18. In that past 12 months have you explicitly used and referenced a reporting guideline checklist in any research report you have published? Yes No I have not published a research paper in the past 12 months I don’t know
19. Which of the following are barriers for you with respect to using reporting guidelines when reporting your research? (Check all that apply)
  a. I don’t know where to find the relevant reporting guideline
  b. I don’t know how to use reporting guidelines
  c. I don’t have time to use reporting guidelines
  d. I don’t see the value in using reporting guidelines
  e. I don’t feel I will get recognition for taking the time to user reporting guidelines
  f. I don’t think my institution prioritizes the use of reporting guidelines
  g. Other, please specify
  **Patient and public Involvement**
20. In that past 12 months have you engaged patients or members of the public in any research you have conducted? Yes No I have not conducted a research project in the past 12 months I don’t know
21. Which of the following are barriers for you with engaging patients or members of the public in your research? (Check all that apply)
  a. I don’t know how to identify patients/public members to contribute
  b. I don’t know how to incorporate patients/public members in my research
  c. I don’t have time to incorporate patients/public members in my research
  d. I don’t see the value in incorporating patients/public members in my research
  e. I don’t feel I will get recognition for taking the time to incorporate patients/public members in my research
  f. I don’t think my institution prioritizes incorporating patients/public members in my research
  g. Other, please specify
22. Is there anything else you want to share? (Free text)

## References

1. https://plus.google.com/+UNESCO. UNESCO Recommendation on Open Science. UNESCO. Published March 2, 2020. Accessed December 17, 2021. https://en.unesco.org/science-sustainable-future/open-science/recommendation

2. Open Science. European Commission - European Commission. Accessed January 7, 2022. https://ec.europa.eu/info/research-and-innovation/strategy/strategy-2020-2024/our-digital-future/open-science_en

3. Government of Canada. Roadmap for Open Science - Science.gc.ca. Accessed September 16, 2020. http://science.gc.ca/eic/site/063.nsf/eng/h_97992.html

4. Open Science - OECD. Accessed January 7, 2022. https://www.oecd.org/sti/inno/open-science.htm

5. G7 Expert Group on Open Science. Open Scholarship Policy Observatory. Published December 1, 2017. Accessed January 7, 2022. https://ospolicyobservatory.uvic.ca/g7-working-group-open-science/

7. Chalmers I, Glasziou P. Avoidable waste in the production and reporting of evidence. The Lancet. 2009;374(9692):786. doi:10.1016/S0140-6736(09)61591-9

8. Baker M. Is there a reproducibility crisis? Nature. 2016;533:452–454. doi:10.1038/533452a

8. Reproducibility in Cancer Biology: Challenges for assessing replicability in preclinical cancer biology | eLife. Accessed January 7, 2022. https://elifesciences.org/articles/67995

10. Anderson JM, Wright B, Rauh S, et al. Evaluation of indicators supporting reproducibility and transparency within cardiology literature. Heart. 2021;107(2):120–126. doi:10.1136/heartjnl-2020-316519

10. Ramirez FD, Motazedian P, Jung RG, et al. Methodological Rigor in Preclinical Cardiovascular Studies: Targets to Enhance Reproducibility and Promote Research Translation. Accessed January 27, 2022. https://www.ahajournals.org/doi/full/10.1161/CIRCRESAHA.117.310628

12. Williams R. Can’t Get No Reproduction. Circ Res. 2015;117(8):667–670. doi:10.1161/CIRCRESAHA.115.307532

13. Begley CG, Ioannidis JPA. Reproducibility in science: Improving the standard for basic and preclinical research. Circ Res. 2015;116(1):116–126. doi:10.1161/CIRCRESAHA.114.303819

14. Bolli R. Reflections on the Irreproducibility of Scientific Papers. Circ Res. 2015;117(8):665–666. doi:10.1161/CIRCRESAHA.115.307496

14. Christensen G, Wang Z, Levy Paluck E, et al. Open Science Practices are on the Rise: The State of Social Science (3S) Survey. Published online January 14, 2020. Accessed January 24, 2022. https://escholarship.org/uc/item/0hx0207r

15. Scherp G, Siegfried D, Biesenbender K, Breuer C. Results report from an online survey among researchers in econo-mics at German higher education institutions in 2019.: 53.

17. Houtkoop BL, Chambers C, Macleod M, Bishop DVM, Nichols TE, Wagenmakers EJ. Data Sharing in Psychology: A Survey on Barriers and Preconditions. Adv Methods Pract Psychol Sci. 2018;1(1):70–85. doi:10.1177/2515245917751886

18. Borghi JA, Gulick AEV. Data management and sharing: Practices and perceptions of psychology researchers. PLOS ONE. 2021;16(5):e0252047. doi:10.1371/journal.pone.0252047

18. Stürmer S, Oeberst A, Trötschel R, Decker O. Early-Career Researchers’ Perceptions of the Prevalence of Questionable Research Practices, Potential Causes, and Open Science. Soc Psychol. Published online November 23, 2017. Accessed March 1, 2022. https://econtent.hogrefe.com/doi/abs/10.1027/1864-9335/a000324

20. Cobey KD, Liu PP. A call to embrace a culture of openness in cardiovascular research. Eur Heart J. Published online May 3, 2022:ehac189. doi:10.1093/eurheartj/ehac189

20. *Open Science Framework*. Accessed June 14, 2016. https://osf.io/

21. Cobey K. An international survey of open science practice in cardiology. Published online September 20, 2022. Accessed May 30, 2023. https://osf.io/v42u8/

23. Baker D, Lidster K, Sottomayor A, Amor S. Two years later: journals are not yet enforcing the ARRIVE guidelines on reporting standards for pre-clinical animal studies. PLoS Biol. 2014;12(1):e1001756. doi:10.1371/journal.pbio.1001756

24. Council AR. ARC Open Access Policy.; 2013. http://www.arc.gov.au/arc-open-access-policy

24. *Tri-Agency Open Access* Policy *on Publications*.; 2017. Accessed November 23, 2016. http://www.cihr-irsc.gc.ca/e/32005.html

26. Following B, Health W, Resolution A, International WHO, Trials C, Platform R. WHO Statement on Public Disclosure of Clinical Trial Results. Published online 2005.

26. ICMJE About ICMJE Clinical Trials Registration. Accessed March 17, 2022. http://www.icmje.org/about-icmje/faqs/clinical-trials-registration/

28. Chan A, Hrobjartsson A, Haahr MT, Gøtzsche PC, Altman DG. Empirical evidence for selective reporting of outcomes in randomized trials. JAMA. 2004;291(20):2457–2465.

29. Fleming PS, Koletsi D, Dwan K, Pandis N. Outcome Discrepancies and Selective Reporting: Impacting the Leading Journals? Plos One. 2015;10(5):e0127495. doi:10.1371/journal.pone.0127495

29. Welcome to Sherpa Romeo - Sherpa Services. Accessed May 19, 2023. https://www.sherpa.ac.uk/romeo/

31. Masum H, Rao A, Good BM, et al. Ten Simple Rules for Cultivating Open Science and Collaborative R&D. PLOS Comput Biol. 2013;9(9):e1003244. doi:10.1371/journal.pcbi.1003244

